# Interferon-*γ* as a Potential Inhibitor of SARS-CoV-2 ORF6 Accessory Protein

**DOI:** 10.1101/2024.01.24.577015

**Authors:** Elena Krachmarova, Peicho Petkov, Elena Lilkova, Dayana Stoynova, Kristina Malinova, Rossitsa Hristova, Anastas Gospodinov, Nevena Ilieva, Genoveva Nacheva, Leandar Litov

## Abstract

ORF6 protein of the SARS-CoV-2 virus plays a crucial role in blocking the innate immune response of the infected cells by inhibiting interferon pathways. Additionally, it binds and immobilises the RAE1 protein onto the cytoplasmic membranes, thereby blocking the transport of mRNA from the nucleus to the cytoplasm. In all these cases the host cell proteins are tethered by the flexible C-terminus of ORF6. A possible strategy to inhibit the biological activity of ORF6 is to bind its C-terminus with suitable ligands. Our in silico experiments suggest that hIFN*γ* binds the ORF6 protein with high affinity, thus impairing its interactions with RAE1 and, consequently, its activity in viral invasion. The here reported in vitro studies reveal a shift of the localization of RAE1 in ORF6 overexpressing cells upon treatment with hIFN*γ* from predominantly cytoplasmic to mainly nuclear, resulting in restoration of the export of mRNA from the nucleus. We also explored the expression of GFP in transfected with ORF6 cells by means of fluorescence microscopy and qRT-PCR, finding that treatment with hIFN*γ* unblocks the mRNA trafficking and reinstates the GFP expression level. The ability of the cytokine to block ORF6 is also reflected in minimising its negative effects on DNA replication by reducing accumulated RNA-DNA hybrids. Our results, therefore, suggest hIFN*γ* as a promising inhibitor of the most toxic SARS-CoV-2 protein.

## 1 Introduction

Coronavirus SARS-CoV-2 caused an unprecedented pandemic that broke out in late 2019. Due to the development of new vaccines and the application of antiviral drugs, the number of cases and deaths have decreased significantly. However, the virus still causes thousands of confirmed new cases per week worldwide [1] with mild to severe symptoms. This data, together with the known asymptomatic expression of SARS-CoV-2 and the emergence of new variants, call for the development of new therapeutic strategies to counteract the virus pathogenicity.

SARS-CoV-2 has a large 30 kb positive sense RNA genome, consisting of 14 open reading frames (ORFs) translated to 29 proteins, including 16 non-structural proteins (NSP1-16), 4 structural proteins, and 9 accessory proteins (ORF3a, 3b, 6, 7a, 7b, 8, 9b, 9c, and 10). The first two groups are essential for the replication, translation, and assembly of the matured viral particles. The accessory proteins play a role in the virus-host protein interactions, autophagy, and apoptosis, and antagonise host immunity [2]. The structural and non-structural proteins of SARS-CoV-2 possess high homology with SARS-CoV proteins. The SARS-CoV-2 accessory proteins show lower homology, the lowest among them belonging to ORF6 (only 69%) [3].

The SARS-CoV-2 ORF6 gene encodes 61 amino acids, of which 40 form its amphipathic N-terminal and 21 — the highly charged C-terminal [4]. Together with some of the other NSPs and accessory proteins (such as NSP6, 13, 14, and 15 and ORF8), ORF6 disrupts innate immune signalling [5], leading to decreased interferon (IFN) signalling in the severe COVID-19 phase [6, 7, 8]. Among all tested SARS-CoV-2 proteins, ORF6 demonstrates the strongest suppression of both primary interferon production and interferon signalling [5]. The ability of ORF6 to suppress the primary immune reaction in the infected cells is manifested in multiple ways. Crucially, it disrupts bidirectional nucleo-cytoplasmic trafficking by interaction with the ribonucleic acid export factor 1 (RAE1) and the nucleopore complex component nucleoporin 98 (NUP98) [9, 10, 11, 12, 13], thus inhibiting host gene expression.

ORF6 was shown to be the most toxic SARS-CoV-2 protein [14]. Recently, we have shown that SARS-CoV-2 ORF6, embedded in the cytoplasmic membranes, binds to RAE1 and sequesters it in the cytoplasm, thus depleting its availability in the nucleus and impairing nucleo-cytoplasmic mRNA transport. This negatively affects the cellular genome stability by compromising the cell cycle progression into the S-phase and by promoting the accumulation of RNA-DNA hybrids, which are a major source of genome instability [9]. In addition, in [15] it was demonstrated that SARS-CoV-2 proteins ORF6 and NSP13 cause degradation of the DNA damage response kinase CHK1 through proteasome and autophagy, respectively. CHK1 loss leads to a deoxynucleoside triphosphate shortage, causing impaired S-phase progression, DNA damage, pro-inflammatory pathways activation and cellular senescence.

All this data prompts an active search for possible inhibitors of SARS-CoV-2 ORF6 activity. To our knowledge, up to now no specific and effective inhibitors of this protein have been reported. Here, we present results from both in silico and in vitro investigations on two potential SARS-CoV-2 ORF6 inhibitors. hIFN*γ* and its C-terminal peptides are shown to be able to form stable complexes with ORF6, thus blocking its activity. Treatment of cells overexpressing ORF6 with hIFN*γ* restores proper subcellular localization of RAE1, thus recovering mRNA transport from the nucleus and leading to a reduction of accumulated RNA-DNA hybrids.

## 2 Materials and Methods

### 2.1 Molecular Modelling

#### 2.1.1 Input Models

The input model for the ORF6 embedded in a model ER membrane was developed previously [9]. The C-terminal hIFN*γ* peptide (aa sequence: LSPAAKTGKRKRSQMLFRGRRASQ) was modeled in a fully stretched configuration using the PyMOL molecular visualization tool [16]. Three separate folding simulations were conducted with different initial velocities, adhering to the molecular dynamics (MD) simulation method outlined in Section 3.1. The combined simulation time was 1.5 *µ*s. The combined trajectory was analyzed through cluster analysis to determine the most frequently occurring conformation of the peptide. For this analysis, the gromos algorithm [17] was employed with a clustering cutoff of 6 Å. The centroid of the largest cluster was used as the input configuration of the CT-hIFN*γ* peptide [18]. The full-length hIFN*γ* model, described in [19] was employed as the input structure for the protein.

#### 2.1.2 MD Protocol

The GROMACS molecular dynamics software [20], version 2021.1 and later was used for the MD simulations. The proteins/peptides were parameterized using the CHARMM36 protein force field [21]. The systems were solvated in rectangular boxes under periodic boundary conditions, ensuring a minimum 2 nm distance of the proteins to the box edges in the Z direction. To neutralize the net charge and ensure physiological salinity, 0.15 mol/l sodium and chlorine ions were added to both systems. The energy of the systems was minimized using the steepest descent method with a maximum force tolerance to 100 kJ/(mol nm). Then a two-stage equilibration was performed with a 50 ps canonical simulation at 310 K, then a 200 ps isothermal-isobaric simulation at the same temperature and 1 atm pressure, utilizing the v-rescale thermostat [22] and Berendsen barostat [23].

The production MD simulations were carried out in the NPT ensemble, maintaining a temperature of 310 K and pressure of 1 atm with the v-rescale thermostat [22] with a coupling constant of 0.25 ps and the Parrinello-Rahman barostat [24, 25] with a coupling constant of 1 ps. The leapfrog integrator [26] was employed with a 2 fs time-step with constraints on bonds between heavy atoms and hydrogens using the PLINCS algorithm [27]. Van der Waals forces were swithed-off from 1.0 nm and cut off at 1.2 nm. Electrostatic interactions were calculated with the smooth PME method [28] using a direct PME cutoff at 1.2 nm.

#### 2.1.3 Synthetic Data Analysis

The MD trajectories were analysed using the standard GROMACS post-processing and analysis tools [29]. The visualisation and manipulation program VMD [30] was used to create all structural figures. The contact maps were generated using the MDTraj package [31] with a heavy-atom cutoff distance of 4.5 Å.

### 2.2 In Vitro Experiments

#### 2.2.1 Cell Culture and Plasmids

WISH (ATCC®CCL-25^™^, ATCC, Manassas, VA, USA) and PC3 (ATCC®CRL-1435^™^) cell lines were grown in MEM and DMEM, respectively, with 10% fetal bovine serum (Gibco^™^, Waltham, MA, USA) and penicillin-streptomycin (10,000 U/mL, Gibco^™^, Waltham, MA, USA). The cells were maintained in a humidified incubator at 37°C with 5% CO2.

The plasmid pLVX-EF1alpha-SARS-CoV-2-orf6-2xStrep-IRES-Puro (Addgene plasmid #141387 from Nevan Krogan) [12] was used for the overexpression of the accessory proteins ORF6. For the overexpression of GFP, the plasmid pEGFP-C1 (NovoPro0#V012024) was used.

The RBD-DsRed plasmid was created by PCR cloning the HB domain of RNAse H1 into the pDsRed-Express-C1 vector (Clontech), following the previously described method [32].

#### 2.2.2 Quantitative Real-Time PCR Analysis

Total RNA from cells, overexpressing ORF6, was extracted by using RNeasy Plus Mini Kit (Qiagen), and 1 *µ*g total RNA from each sample was reverse-transcribed using RevertAid H Minus First Strand cDNA Synthesis Kit (Thermo Scientific^™^) according to the kit instructions. For isolation and purification of the nuclear RNA, the Cytoplasmic and Nuclear RNA Purification Kit (Norgen biotek corp.) was used in accordance with the manufacturer’s instructions.

Relative expression levels of the target GFP gene was assessed by qRT-PCR analysis using the SYBR^™^ Select Master Mix (Thermo Scientific^™^). The *β*-actin housekeeping gene was used as an internal control for the normalisation of gene expression.

The sequences of the primer oligonucleotide of the studied genes are listed in Suppl. Table S1. The analysis was performed on Rotor-Gene 6000 thermal cycler (Corbett, QIAGEN, Hilden, Germany). The gene expression data were analysed by using Rotor-Gene 6000 Software (QIAGEN). The relative expression levels of the target genes were normalised to the endogenous control specific to each sample. Each qRT-PCR reaction was carried out with a minimum of three replicates across distinct PCR runs. Statistical significance was determined using a t-test, and significance was denoted by values less than 0.05.

#### 2.2.3 Immunofluorescence Microscopy

WISH and PC3 cells were cultured on 12 mm coverslips (Epredia) and transfected with plasmids, relevant to the experiment being performed. Cells were fixed with 3,7% formaldehyde in 1 × PBS for 10 min at room temperature, followed by treatment with methanol for 10 min at −20°C and permeabilisation with 0,5% Triton X-100 in PBS for 5 min. Coverslips were then blocked for 1 h in blocking buffer (5% BSA and 0,1% Tween 20 in × 1 PBS) and incubated overnight at 4°C with primary antibody (in blocking buffer). After washes cells were stained for 1 h with a secondary antibody at room temperature.

The nuclei were stained with DAPI (Cell Signalling Technology) and the coverslips were washed and mounted using ProLong^™^ Gold Antifade mounting media (Invitrogen). Fluorescent images were obtained by using Zeiss Axiovert 200M fluorescence inverted microscope and analysed by CellProfiler software [33].

#### 2.2.4 hIFN*γ* Treatment

Following transfection, cells were treated with 100 ng/ml hIFN*γ* purified as described in [34] for 24 h 37°C and 5% CO2. Depending on the methodology being performed, cells were either fixed or collected by trypsinisation for further analysis.

#### 2.2.5 Fluorescence Recovery After Photobleaching (FRAP) Analysis

Cells were transfected with the RBD-DsRed plasmid expressing the fusion between the RNA Binding Domain of RNAse H1 and DsRered. Andor Revolution XDI spinning disk confocal system was used to perform the FRAP analysis. For the duration of the experiment, cells were maintained in a heated chamber in CO2-independent medium. Using a bleaching pulse applied at the fifth second, images were taken every 1 s for 150 s. For measuring the intensity and further analyses, CellTool [35] and EasyFrap software [36] were used.

#### 2.2.6 DNA Fiber Labelling

DNA fibre analyses were carried out in accordance with standard procedure [MM7] with slight adjustments. In brief, PC3 cells that were growing exponentially were first incubated for 10 minutes at 37°CC and 5% CO2 with 25 *µ*M chlorodeoxyuridine (CldU) and then with 250 *µ*M iododeoxyuridine (IdU) at the same conditions. Spreads were made using 2500 cells that were suspended at 1×106 cells/ml in 1× PBS. Fibre lysis buffer (50 mM EDTA and 0.5% SDS in 200 mM Tris-HCl, pH 7.5) was used to lyse the cells. To disperse DNA fibres, the slides were tilted by about 25 degrees until the fibre solution drop reached the bottom of the slide, then was allowed to dry. After drying, the slides were immersed in 2.5 M HCl for 80 minutes, rinsed with PBS and blocked in 5% BSA in 1xPBS for 40 minutes. Primary antibodies were diluted in blocking buffer and applied overnight: rat anti-BrdU antibody (Abcam cat # Ab6326) was used to detect CldU, and mouse anti-BrdU antibody (Becton Dickinson, cat # 347580) was used to detect IdU. After washing, slides were incubated for 60 minutes with secondary antibodies, goat anti-mouse DyLight®488 (Abcam, 96879) and goat anti-rat DyLight®594 (Abcam, 96889). ProLong Gold anti-fade reagent (Invitrogene) was used to mount the slides. Images were taken using a Carl Zeiss Axiovert 200M microscope that was outfitted with an Axiocam MR3 camera. Measurements of fibre length were performed using Image J or the DNA size finder software [37].

#### 2.2.7 Statistical Analysis

The in vitro data was collected by at least three independent measurements of each data point. As presented in Section 2, the experimental numbers and figures are based on the mean value ± the standard deviation. Statistical significance was estimated using Student’s t-test for independent pairs.

## 3 Results

### 3.1 Molecular Modelling of the Interaction of SARS-CoV-2 ORF6 and hIFN*γ*/hIFN*γ* C-terminal Peptides

As discussed in [9], experimental data indicates that the C-terminal domain of the SARS-CoV-2 ORF6 protein is responsible for its biological activity [11, 38]. The ORF6 C-terminus is a solvent-exposed highly negatively charged flexible tail [9]. It is to be expected that it will interact with high affinity with a positively charged peptide/protein and that this non-specific interaction will be able to inhibit the binding of ORF6 to the host export factor RAE1. We propose the C-terminal peptide of the cytokine hIFN*γ* (CT-hIFN*γ*) as a suitable candidate for SARS-CoV-2 ORF6 inhibitor. CT-hIFN*γ* is a 21 residues long flexible non-structured tail, containing two domains rich in basic amino acids — D_1_ (^125^KTGKRKR^131^) and D_2_ (^137^RGRR^140^). The sequences of the two C-terminal segments are listed in Table 1.

**Table 1.**
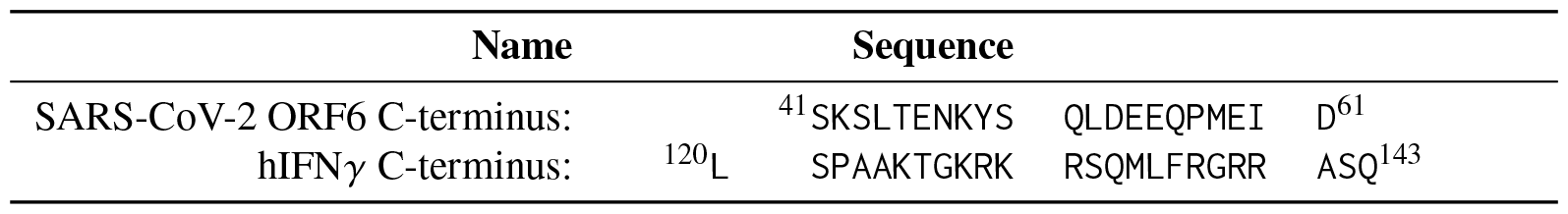
Sequences of the C-terminal tails of the ORF6 and hIFN*γ* proteins.

To study the ORF6 interaction with the CT-hIFN*γ* peptide, a molecular dynamics (MD) simulation was set up, where three CT-hIFN*γ* molecules were placed at a distance of about 3–5 nm above a single ORF6 protein, embedded in a 7.5 × 7.5 nm^2^ model endoplasmic reticulum (ER) membrane. The input configuration is displayed in Suppl. Fig. S1. The total simulation time was 280 ns.

The CT-hIFN*γ* peptides were immediately electrostatically attracted to ORF6 and began to move towards the membrane. Within the first few nanoseconds, the viral protein formed numerous close contacts with one of the peptides, CT-hIFN*γ*-1. This complex underwent some readjustment after the initial intense interaction. The number of close contacts changed slightly from 64 ± 27 in the first half to 54 ± 24 in the second half of the simulation (Suppl. Fig. S2a), while the few H-bonds that were formed between CT-hIFN*γ*-1 and ORF6 dropped from 2.5 ± 1.3in the beginning to 1.3 ± 0.7 in the second half of the simulation (Fig. 1a).

**Figure 1.**
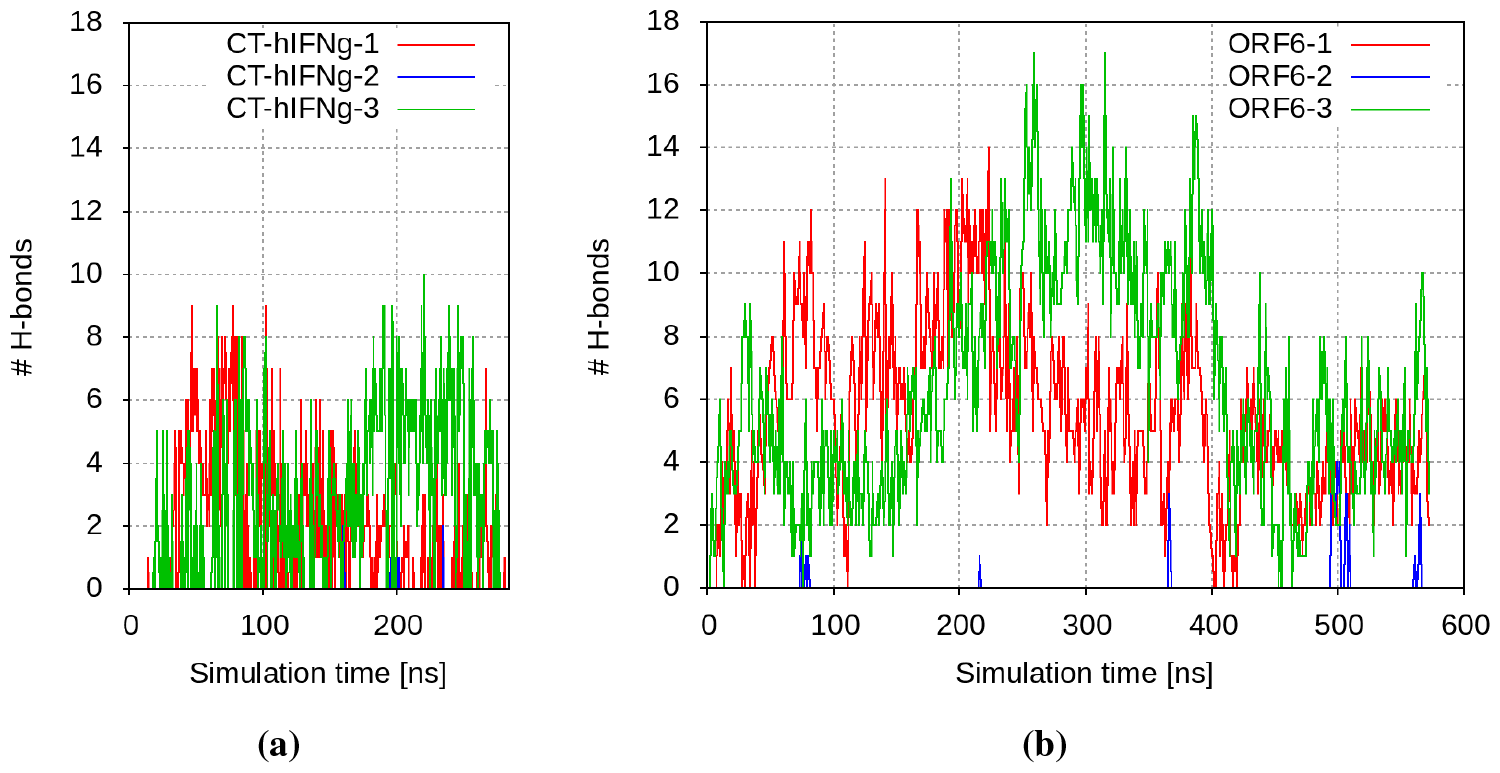
Number of H-bonds between (a) ORF6 and the CT-hIFN*γ* peptides; and (b) hIFN*γ* and the ORF6 proteins.

Right after CT-hIFN*γ*-1, ORF6 bonded to a second peptide as well, CT-hIFN*γ*-3. On average this complex was maintained by 66 ± 20 close contacts and 3.6 ± 1.3 H-bonds in the second half of the simulation (Suppl. Fig. S2a and Fig. 1a, respectively).

The ORF6 protein also attracted the third CT-hIFN*γ* peptide and formed some transient contacts with it, but this complex was not stable. This was probably due to the electrostatic repulsion of this molecule from the other two bonded peptides.

The contact maps between the ORF6 protein and the two binding C-terminal hIFN*γ* peptides are shown in Fig. 2a. The permanent contacts in the case of CT-hIFN*γ*-1 involved residues ^56^QPMEID^61^ of ORF6 and residues ^132^SQML^135^ of the peptide, while the CT-hIFN*γ*-3 peptide interacted via the somewhat extended domain ^131^RSQMLF^136^, as well as the basic residues R^129^ and R^140^ of its D_1_ and D_2_ domains with the longer negatively charged sequence ^52^LDEEQPM^58^ and I^60^ of ORF6 (right panel of Fig. 2a). In both cases, the C-terminal peptides remained in a stable complex with the viral protein.

**Figure 2.**
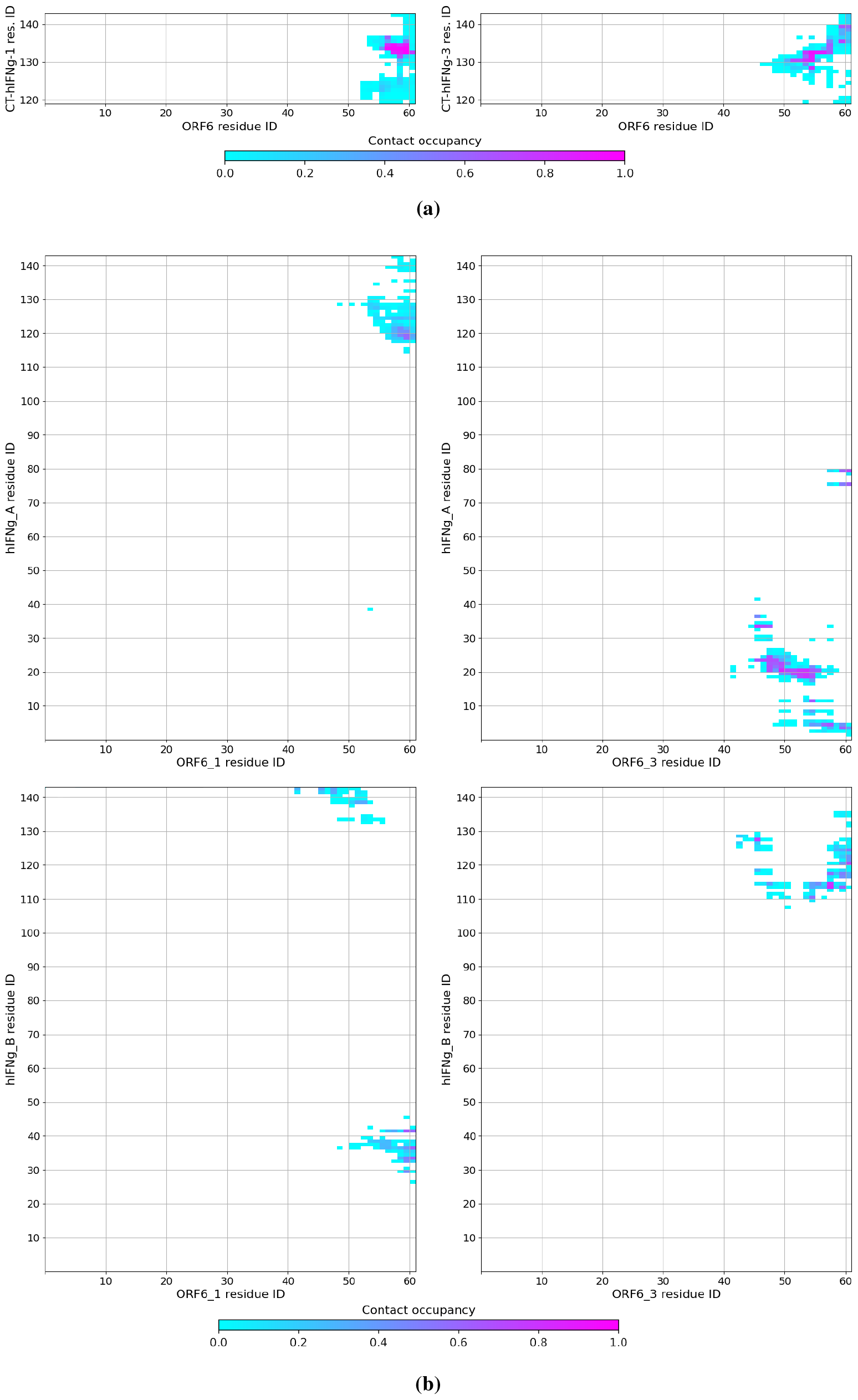
Maps of the close contacts between (a) ORF6 and the CT-hIFN*γ* peptides; and (b) hIFN*γ* and the ORF6 proteins.

Based on these results, a second MD simulation was carried out to probe the interaction of the whole hIFN*γ* cytokine with the SARS-CoV-2 proteins. The second system consisted of four ORF6 proteins embedded in a 15 ×15 nm^2^ membrane and one full-length hIFN*γ* homodimer placed close to the membrane between the C-termini of the four ORF6 molecules at a distance of about 4.5–5 nm. The initial configuration of this simulation is presented in Suppl. Fig. S3.

The simulation confirmed the expected rapid binding process: two of the four ORF6 molecules became engaged in close contacts and started forming H-bonds with the cytokine already within the first 20-50 ns, as seen in Fig. 1b and Fig. 2b. However, besides its C-termini (amino acid residues ^119^ELSP^122^, K^125, 128^KR^129^, R^139, 142^SQ^143^), this process also involved solvent-exposed parts of the globule, containing polar or charged amino acids — ^4^YV^5^, K^12, 19^HSDVVADN^25^, L^30, 33^LK^34^, K^37^, R^42^, D^76^, K^80^, H^1^11, ^114^IQ^115^, A^118^. The complexes so-formed were stable and maintained by numerous contacts and several H-bonds. A typical conformation of the full-length hIFN*γ* homodimer bound to three of the four ORF6 proteins is shown in Fig. 3.

**Figure 3.**
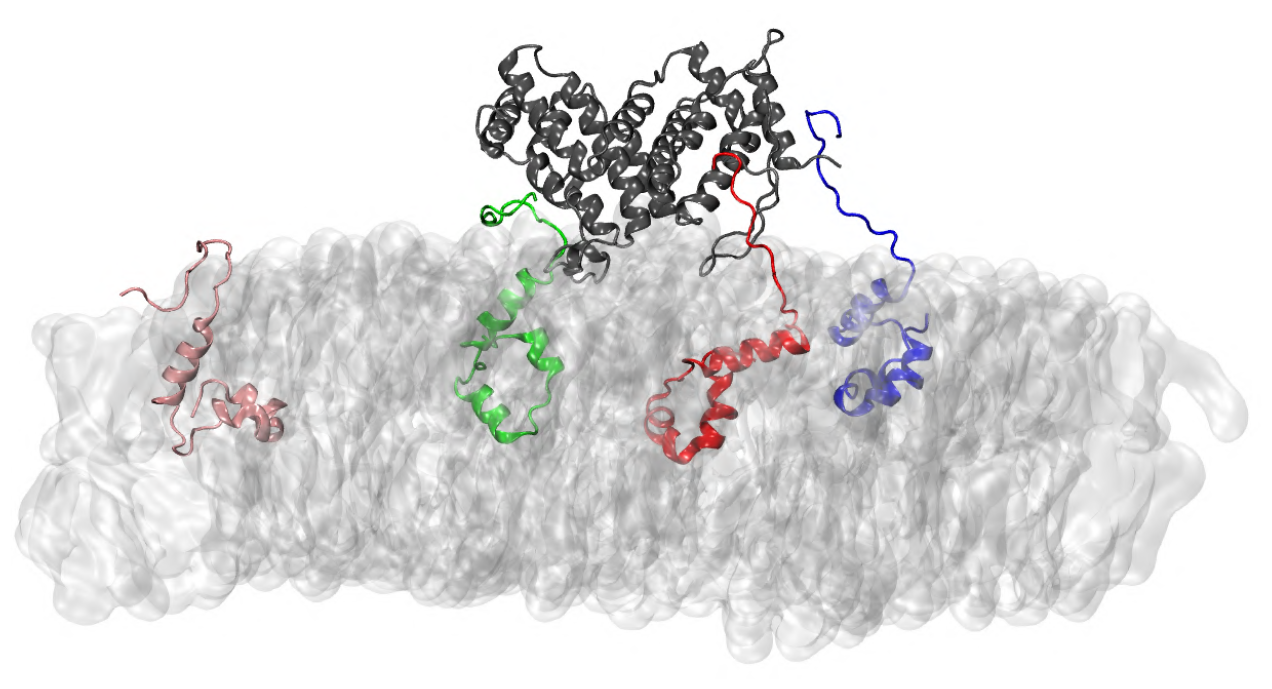
Binding of hIFN*γ* and three ORF6 proteins. hIFN*γ* is depicted in dark gray, ORF6-1 to ORF6-4 are shown respectively in red, blue, green and pink cartoons. The ER membrane is in light gray surface representation.

The results of our in silico experiments suggest that hIFN*γ* binds the ORF6 protein with high affinity, impairing its interactions with RAE1 and thus inhibiting its activity in viral invasion.

### 3.2 hIFN*γ* Restores the Cellular Localization of Rae1

To experimentally test this hypothesis, we drew on our earlier result [9] that the co-localization of ORF6 and one of its main targets, the mRNA export factor Rae1 [39], leads to impairment of mRNA transportation due to shifting RAE1’s cellular localisation from mainly nuclear to predominantly cytoplasmic [9]. By means of immunofluorescence analysis we studied the changes in the in-cell localisation of RAE1 in WISH cells overexpressing ORF6 upon treatment with hIFN*γ*. As seen in Fig. 4a, the co-localization of RAE1 and ORF6 is hindered in the hIFN*γ*-treated cells. The cytokine restores the nuclear localisation of RAE1 (Fig. 4b), most probably resulting in unblocking of mRNA transportation.

**Figure 4.**
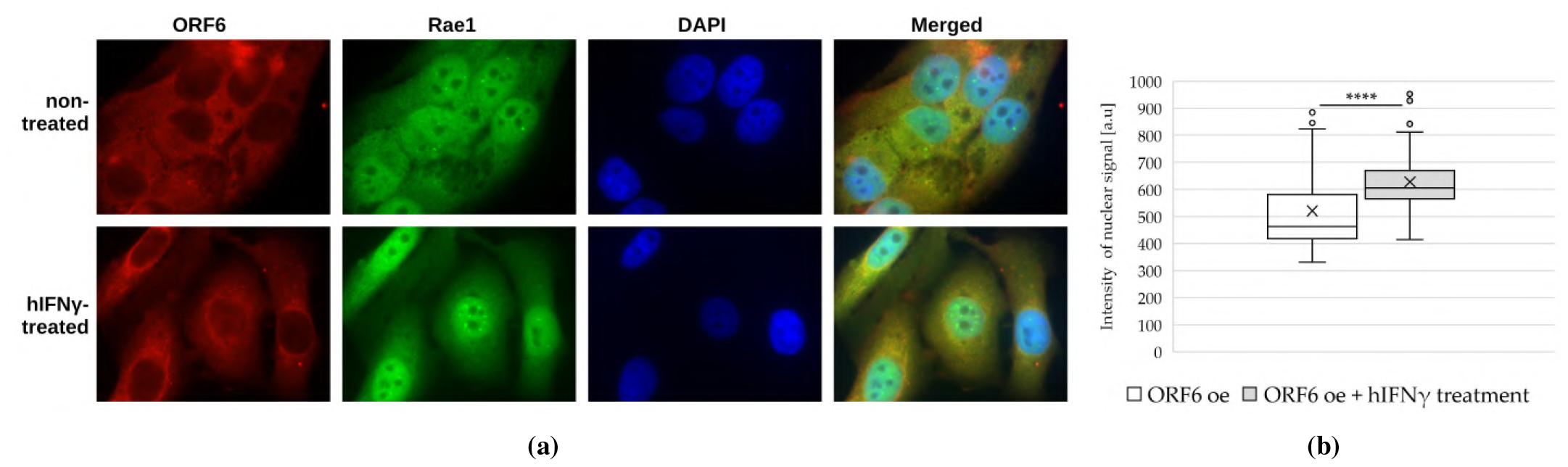
RAE1 localization in ORF6 overexpressing cells upon hIFN*γ* treatment. (a) Non-treated and hIFN*γ*-treated cells, overexpressing ORF6, were stained with antibodies against RAE1 and ORF6. Representative images are shown. (b) The fluorescence intensity of RAE1 (in the green channel) was analysed for each cell using the ImageJ software. The difference is statistically significant with a ^****^ p-value < 0.0001, estimated using Student’s test for three independent experiments (two-tailed unpaired Student’s test).

Further, we monitored the RAE1 mRNA levels in ORF6 overexpressing cells treated with hIFN*γ*. For this purpose, we performed qRT-PCR of total RNA, isolated from transfected WISH cells, both non-treated and treated with hIFN*γ*. The data analysis presented in Fig. 5 shows that the treatment with the cytokine leads to elevated RAE1 mRNA levels in cells overexpressing ORF6.

**Figure 5.**
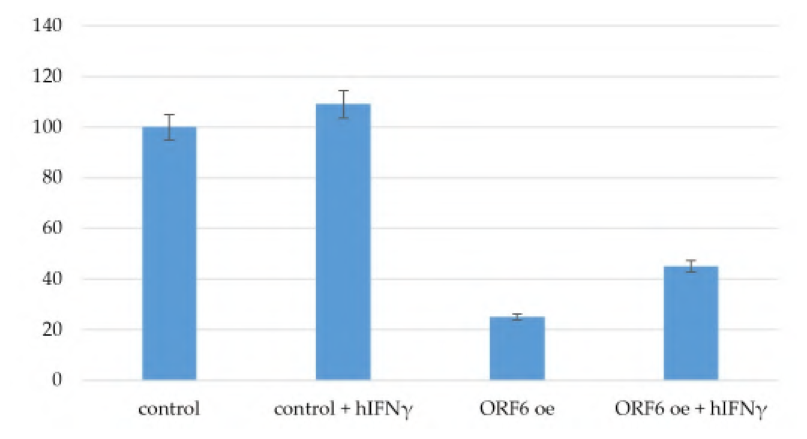
Levels of expression of RAE1 in ORF6 overexpressing cells upon hIFN*γ* treatment. Control – cells, transfected with empty vector; control + hIFN*γ* – cells transfected with empty vector and treated with hIFN*γ*; ORF6 oe – cells transfected with plasmid coding for ORF6; ORF6 oe + hIFN*γ* – cells transfected with plasmid coding for ORF6 and treated with hIFN*γ*. The figures shown are based on three independent experiments and are represented as mean ± standard error of the mean (error bars).

The observed downregulation of RAE1 expression in transfected cells is most likely due to a general inhibition of mRNAs export from the nucleus, leading to the suppression of a number of processes in the cell, including RAE1 expression. The increase in RAE1 expression level after hIFN*γ* treatment could be explained by the overall unblocking of mRNA export due to the hIFN*γ* inhibition of the ORF6/RAE1 interaction, which leads to RAE1 sequestration in the cytoplasm.

### 3.3 Effect of hIFN*γ* Treatment on GFP Fluorescence Intensity

To further investigate the role of hIFN*γ* as a potential inhibitor of ORF6, we established a model system in which we used the reporter GFP protein to monitor the fluorescence intensity upon ORF6 overexpression. For this reason, we co-transfected PC3 cells with plasmids coding for ORF6 and GFP and analysed the overall intensity of the GFP fluorescence after hIFN*γ* treatment.

As seen from the fluorescence images (Fig. 6a) and their statistical analysis (Fig. 6b), the overexpression of ORF6 significantly reduces the overall GFP fluorescence intensity levels, confirming the negative effect of ORF6 on the mRNA export due to the depletion of the nuclear RAE1. The treatment with hIFN*γ*, however, restored the overall fluorescence signal, confirming our hypothesis on the role of hIFN*γ* as an ORF6 inhibitor.

**Figure 6.**
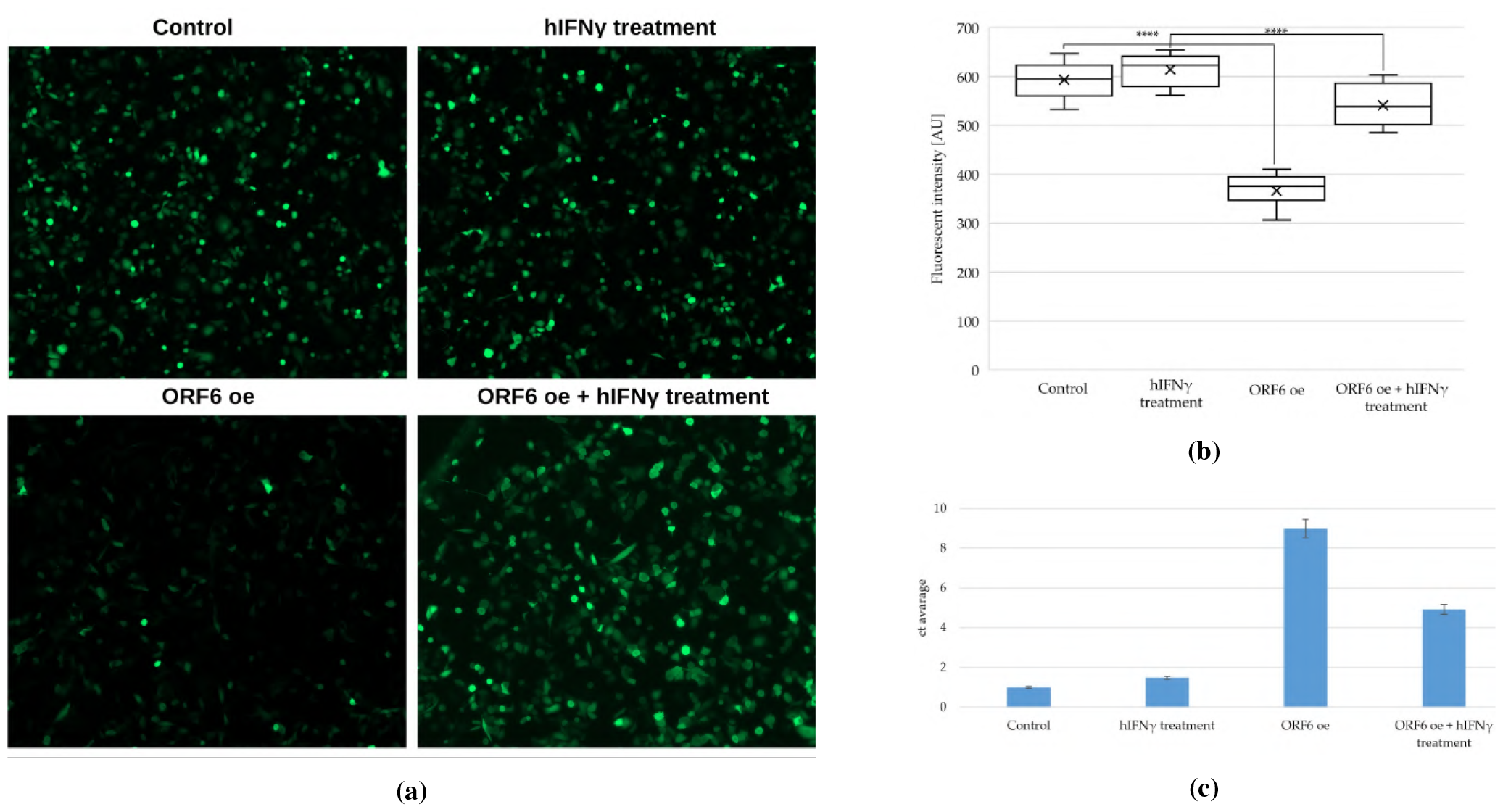
GFP reporter expression. (a) Visualisation of GFP expression under fluorescence microscope. Representative images are shown. (b) The fluorescence intensity of GFP was analysed using ImageJ software: ^****^ p-value < 0.0001, based on three independent experiments (two-tailed unpaired Student’s t-test) (c) qRT-PCR of the nuclear levels of GFP mRNA isolated from (a). Control – cells, transfected with GFP; hIFN*γ* treatment – cells transfected with GFP and treated with hIFN*γ*; ORF6 oe – cells co-transfected with plasmids coding for ORF6 and GFP; ORF6 oe + hIFN*γ* – cells co-transfected with plasmids coding for ORF6 and GFP and treated with hIFN*γ*. ^****^ p-value < 0.0001, based on three independent experiments (two-tailed unpaired Student’s t-test).

Further, we analysed the levels of nuclear GFP mRNA upon hIFN*γ* treatment. For this reason, we performed qRT-PCR of the nuclear mRNA fraction isolated from PC3 cells, co-expressing ORF6 and GFP and treated with hIFN*γ*. The data analysis revealed that the nuclear accumulation of mRNA in the transfected PC3 cells following hIFN*γ* treatment is significantly reduced compared to the levels in transfected non-treated cells (Fig. 6c). These observations clearly show the ability of hIFN*γ* to inhibit the ORF6-induced obstruction of mRNA transport and, as a result, to restore protein expression levels.

### 3.4 hIFN*γ* Prevents R-loop Formation and Restores the Replication Fork Rates

We have previously shown that the sequestration of RAE1 in the cytoplasm by ORF6 impedes the mRNA transportation and leads to accumulation of R-loops, thus hindering the progression of active replication forks and inducing genome instability [9]. To study whether the cellular proliferation defects caused by ORF6 can be reverted by hIFN*γ* treatment, we performed DNA fiber labelling analysis of PC3 cells overexpressing ORF6 and treated with hIFN*γ* by measuring the lengths of second label (green) segments of red-green tracks. The obtained data showed that hIFN*γ* effectively countered the detrimental effects of the stalled mRNA export caused by ORF6 and restored replication fork rates to the levels of the control cells (Fig. 7a). We further performed FRAP analysis on living cells following the mobility of the RNA binding domain of RNAse H1 fused to DsRed (RBD–DsRed). The results (Fig. 7b) indicated that the rate of fluorescence recovery was restored in cells expressing ORF6 and treated with hIFN*γ*. The restoration in the mobile fraction of RBD–DsRed observed in these cells indicated that they accumulate fewer R-loops than the cells expressing ORF6.

**Figure 7.**
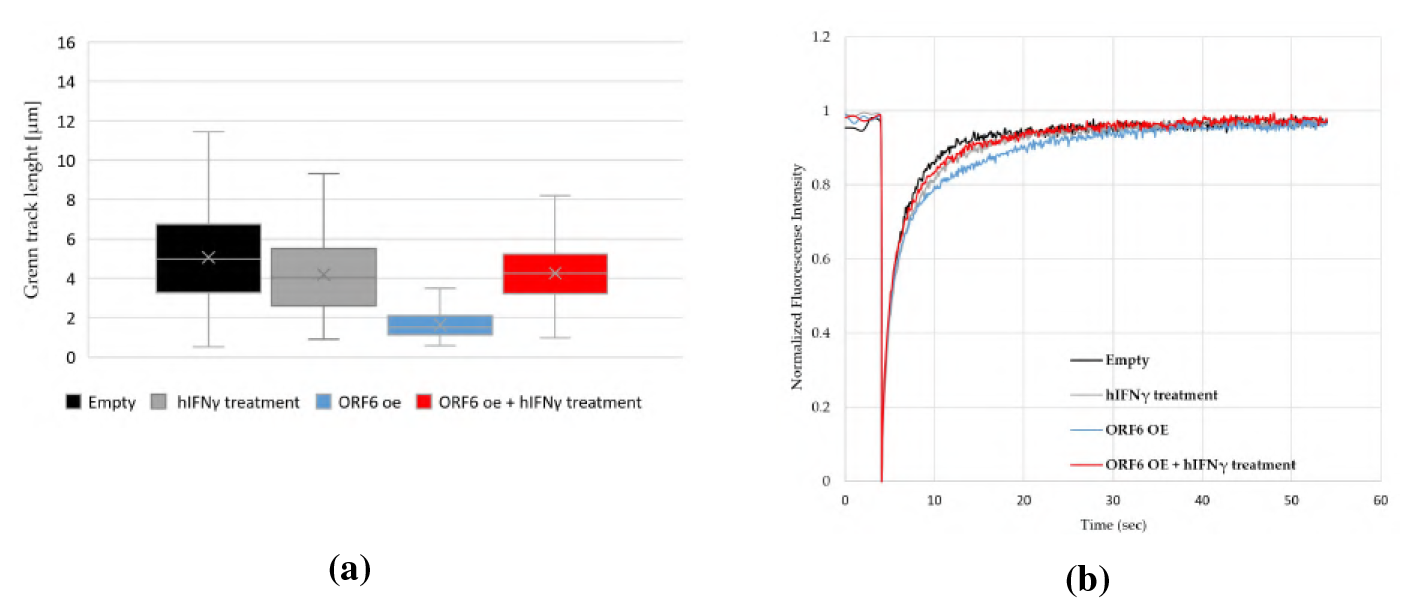
hIFN*γ* inhibits the formation of R-loops and reinstates the rates of replication fork progression. (a) Replication rates cells, ^****^p-value < 0.0001, based on three independent experiments (two-tailed unpaired Student’s t-test). (b) FRAP analysis of RBD-DsRed in control, ORF6-overexpressing cells, and ORF6-overexpressing cells, treated with hIFN*γ*.

## 4 Discussion

ORF6 is one of the most toxic proteins of the SARS-CoV-2 virus and contributes significantly to the viral pathogenicity [40, 4, 14]. It has recently been shown, using a broadly applicable live-cell dose-response pipeline, that SARS-CoV-2 ORF6 is about 15 times more potent than its orthologue from SARS-CoV lineages as a type I interferon pathways antagonist [41].

ORF6 affects in several different ways the innate immune reaction of the cell by blocking the IFN induced pathways and mRNA transportation from the nucleus to the cytoplasm. The interaction of ORF6 with the nucleopore complex RAE1-NUP98 and importin *α*/importin*β*1 blocks the necessary for the activation of interferon-inducible genes translocation of the transcription factors STAT1, STAT2 and IRF3 into the nucleus [10, 42]. Moreover, it was shown that through its C-terminus ORF6 directly binds STAT1 independently of IFN*γ* stimulation [43]. Recently, it has also been shown that ORF6 inhibits the TNF-*α* induced NF-_*K*_B activation by suppressing p65 translocation into the nucleus, most probably mediated by an interaction with importin *α*1 [44]. Using high-content screening and computational prediction, it was shown that together with only two other SARS-CoV-2 proteins ORF6 inhibits all three IFN signalling pathways [45].

Experimental data shows that in infected cells ORF6 localises on cytoplasmic membranes (endoplasmic reticulum, Golgi apparatus, autophagosomes, and lysosomes) [46, 14, 47] where it binds RAE1 with high affinity and thus blocks mRNA export from the nuclei. We have recently shown that the N-terminal region of ORF6 is incorporated into the membrane and the highly negatively charged C-terminus interacts with RAE1 [9].

The ORF6-RAE1 interaction leads to inhibition of mRNAs encoding of key antiviral factors, like IFN-*β*, IL-6, IRF1 (antiviral transcription factor), RIG-I (cytosolic pattern recognition receptor responsible for the type I interferon response) and ZNFX1 (RNA-binding protein that is required to restrict the replication of RNA viruses by interaction with Mitochondrial Activation of Viral Signalling (MAVS) to initiate the type I interferon response), that are both basally expressed and induced by type I interferons [48, 49]. In addition, a direct interaction of ORF6 with MAVS is observed, associated with inhibited RIG-I 2CARD-mediated IFNB1 promoter activation [50].

The common feature in all interactions of ORF6 with host cell proteins is that all these proteins bind to the flexible C-terminus of ORF6 and become immobilized on the cytoplasmic membranes. A potential strategy to inhibit the biological activity of ORF6 is to bind its C-terminus with a suitable ligand. Here we propose as such both the full-length hIFN*γ* homodimer and its highly positively charged unstructured C-terminal peptides.

hIFN*γ* is the only representative of type II interferons. It is a basic (alkaline) protein consisting of 143 amino acids (aa), 28 of which are Lys and Arg. They are organised in six *α*-helices comprising 62% of the molecule, connected by unstructured regions [51], and highly positively charged 21 aa long C-terminus. The active form of hIFN*γ* is a very stable non-covalent homodimer. The latter activates the target cells via interaction with the extracellular (called “soluble”) domain of hIFN*γ* receptor [52] followed by activation of the JAK/STAT1 transduction pathway [53].

hIFN*γ* is endowed with multiple biological effects through orchestration of a broad spectrum of distinct cellular programs. Despite the extraordinary complexity of the hIFN*γ* response, the major function that can be attributed to this cytokine is regulation of the innate and adaptive immunity. The properties of hIFN*γ* include also: stimulation of antiviral and bactericidal activity, increase of antigen presentation, activation and increase of lysosome activity in macrophages, suppression of Th2 cell activity, promotion of adhesion and binding of leukocytes (required for leukocyte migration), promotion of Th1 differentiation by upregulating the transcription factor T-bet, promotion of NK cell activity, and more general effects on cell proliferation and apoptosis [54, 55].

Our in silico studies show that both the C-terminus and the whole hIFN*γ* molecule are able to form stable non-covalent complexes with the SARS-CoV-2 ORF6 protein. In both cases, the interaction engages the C-terminus of ORF6, particularly the region that is responsible for binding of host proteins [11, 38]. This could potentially render the cytokine and/or its C-terminal peptide suitable competitors for the viral protein, preventing, in particular, its binding to and sequestration of RAE1 onto cytoplasmic membranes. This hypothesis is indirectly supported by our experimental data — the treatment with hIFN*γ* restores the nuclear localisation of RAE1, thus suggesting that the cytokine blocks the ability of ORF6 to interact with RAE1. An indication that ORF6 is blocked by hIFN*γ* and the mRNA transport is restored lies in the observed elevated expression of RAE1 and GFP.

For the hIFN*γ* to compete effectively with RAE1 for the ORF6 protein, the formation of the ORF6-hIFN*γ* complex should be thermodynamically more favourable than the formation of the ORF6-RAE1 complex, that is, the overall Gibbs free energy change (Δ*G*_*bind*_) for the former should be lower. Binding free energy results from electrostatic interactions, hydrophobic effect, and hydrogen-bonds formation. The entropic term of Δ*G*_*bind*_ due to the hydrophobic effect is proportional to the change in the solvent-accessible surface area of the hydrophobic residues in the C-terminus of ORF6 upon binding [56]. However, for hIFN*γ* and RAE1 the value is virtually the same (Suppl. Fig. S4, data based on previous results [9]). Therefore, the difference in ORF6 binding free energy to hIFN*γ* and RAE1 should be determined by the difference in the ionic / H-bond interactions. On average, RAE1 forms 20% more H-bonds with the ORF6 C-termini than the full-length hIFN*γ* homodimer — 18 vs. 15. On the other hand, three times more basic amino acids from the cytokine are involved in the binding than from the mRNA transport protein — 9 vs. 3. Thus, the difference between RAE1 and hIFN*γ* interactions with ORF6 is predominantly determined by the electrostatic interactions. Additionally, at large intermolecular distances, the long-range electrostatic interactions favourably influence binding affinity and provide a substantial enthalpic contribution to the stability of the complex. Hence, the hIFN*γ* homodimer with its net positive charge of +18e has a significant advantage over the neutral RAE1 protein.

We explored the expression of GFP in cells transfected with ORF6 by fluorescence microscopy and qRT-PCR using a similar reporter system to the one in [57]. The obtained results undoubtedly showed that treatment with hIFN*γ* unblocked the mRNA trafficking, as it reinstated the GFP expression level.

The obstruction of mRNA export has been shown to lead to the accumulation of R-loops [58]. Our previous results showed that the overexpression of ORF6 causes inhibition of cell cycle and accumulation of R-loops, thus affecting cellular proliferation and inducing genome instability [9]. By means of fiber labelling analysis we demonstrated that hIFN*γ* was able to revert the DNA replication forks’ progression. Through FRAP experiments in living cells, we observed restored mobility of RBD–DsRed in ORF6-expressing cells upon treatment with the cytokine.

Our in vitro data presented here show that hIFN*γ* is able to effectively block the activity of ORF6, which points to hIFN*γ* as a potential inhibitor of this SARS-CoV-2 accessory protein.

As already mentioned, SARS-CoV-2 infection disrupts innate immune signalling [4] leading to decreased IFN signalling in the severe COVID-19 phase [55]. On the basis of available data on the pathology of MERS-CoV, SARS-CoV, and SARS-CoV-2, a synergistic use of type I and type II IFNs has been suggested as therapeutic means against COVID-19 [55]. Treatment with interferons is proposed in order to boost the immune system to be able to counteract the viral propagation. Here we propose a completely different mechanism by which hIFN*γ* is able to neutralise the negative effects of SARS-CoV-2 infection, i.e. by suppressing the activity of ORF6, the most toxic SARS-CoV-2 accessory protein. However, hIFN*γ* plays a dual role in the immune system — immunomodulatory and proinflammatory. During different viral infections, including SARS-CoV-2, the cytokine was reported to manifest both effects, mainly depending on the stage of the disease [59]. Therefore, timing, duration and dosage of application would be critical for the result of the treatment. In this context, application of the C-terminal peptides of hIFN*γ* alone might be the winning strategy. Further research is necessary to explore this in detail.

## 5 Conclusion

Here we report an in silico and in vitro evaluation of hIFN*γ* and its C-terminal peptide as potential inhibitors of the SARS-CoV-2 ORF6 protein. Our simulations demonstrate that both the cytokine and its C-termini effectively bind the C-terminal part of ORF6, thus blocking it and preventing the viral protein from binding RAE1. The conducted in vitro assays show that, indeed, after treatment with hIFN*γ* the localization of RAE1 in ORF6 overexpressing cells is shifted from predominantly cytoplasmic to mainly nuclear. As a result, the export of mRNA from the nucleus is restored. The ability of this cytokine to block ORF6 is also shown by the recovery of its negative effects on DNA replication, i.e. by reduction of accumulated RNA-DNA hybrids. Our data puts forward hIFN*γ* as a promising inhibitor of ORF6, one of the most toxic SARS-CoV-2 proteins.

## Acknowledgments

WeThis research was partially funded by the Bulgarian National Science Fund under Grant KP-06-DK1/5/2021 SARSIMM.

Computational resources were provided by the Discoverer supercomputer thanks to Discoverer PetaSC and EuroHPC JU, as well as the BioSim HPC cluster at the Faculty of Physics, Sofia University “St. Kliment Ohridski”.

## Abbreviations

The following abbreviations are used in this manuscript:

aa: amino acids
ACTB: actin beta
CHK1: checkpoint kinase 1
CT-hIFN*γ*: C-terminal human interferon-*γ* peptide
EFGP: green fluorescent protein
ER: endoplasmic reticulum
FRAP: fluorescence recovery after photobleaching
GFP: green fluorescent protein
hIFN*γ*: human interferon-*γ*
MD: molecular dynamics
NSPs: non-structural proteins
NUP98: nucleoporin 98
ORF6 oe: ORF6-overexpressing cells ORF6 open reading frame 6
ORFs: open reading frames
qRT-PCR: real time quantitative PCR
RAE1: ribonucleic acid export 1
SASA: solvent accessible surface area

## Supplementary Material

**Figure S1.**
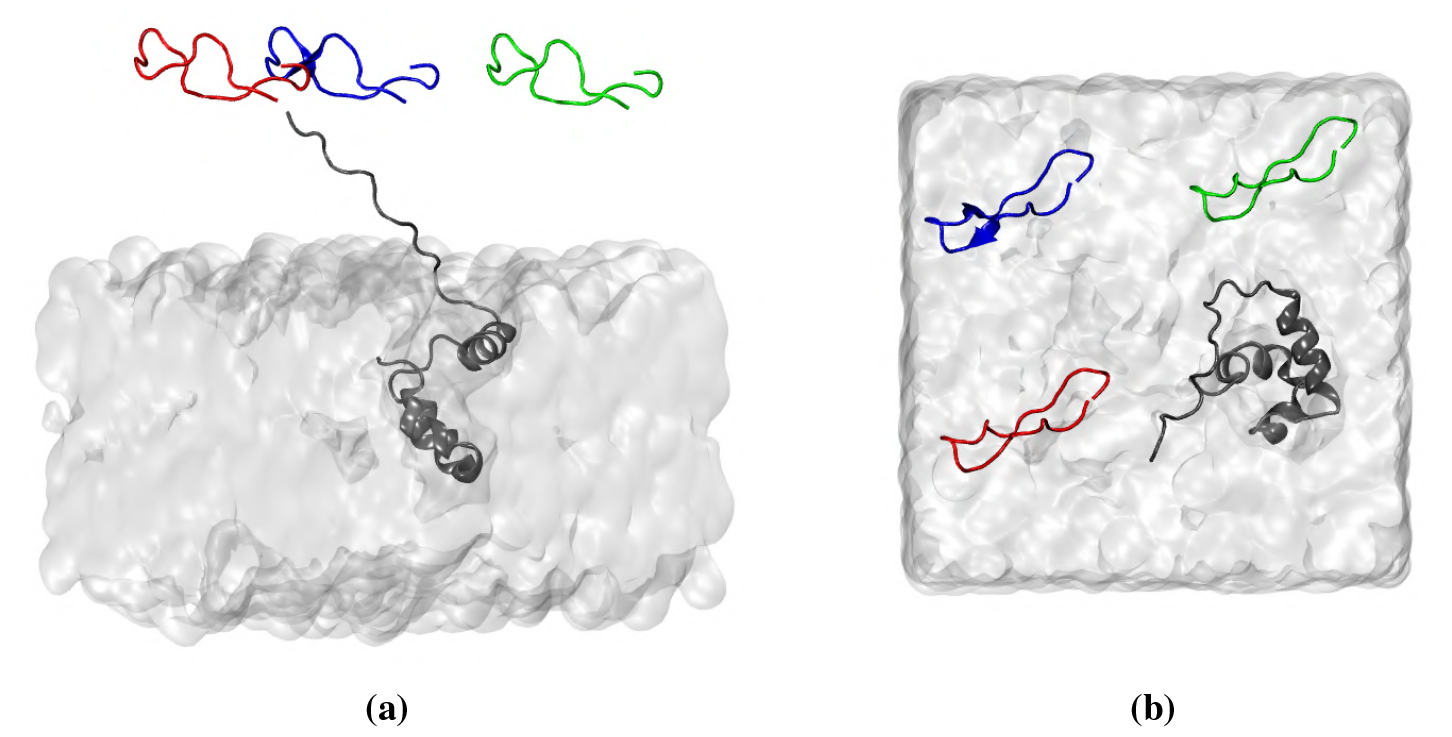
(a side view and (b) top vie of the input configuration of an ORF6 protein, embedded in a model ER membrane and three CT-hIFN*γ* peptides. ORF6 is depicted in dark gray, CT-hIFN*γ*-1 to CT-hIFN*γ*-3 are shown respectively in red, blue, and green. The ER membrane is in light gray surface representation.

**Figure S2.**
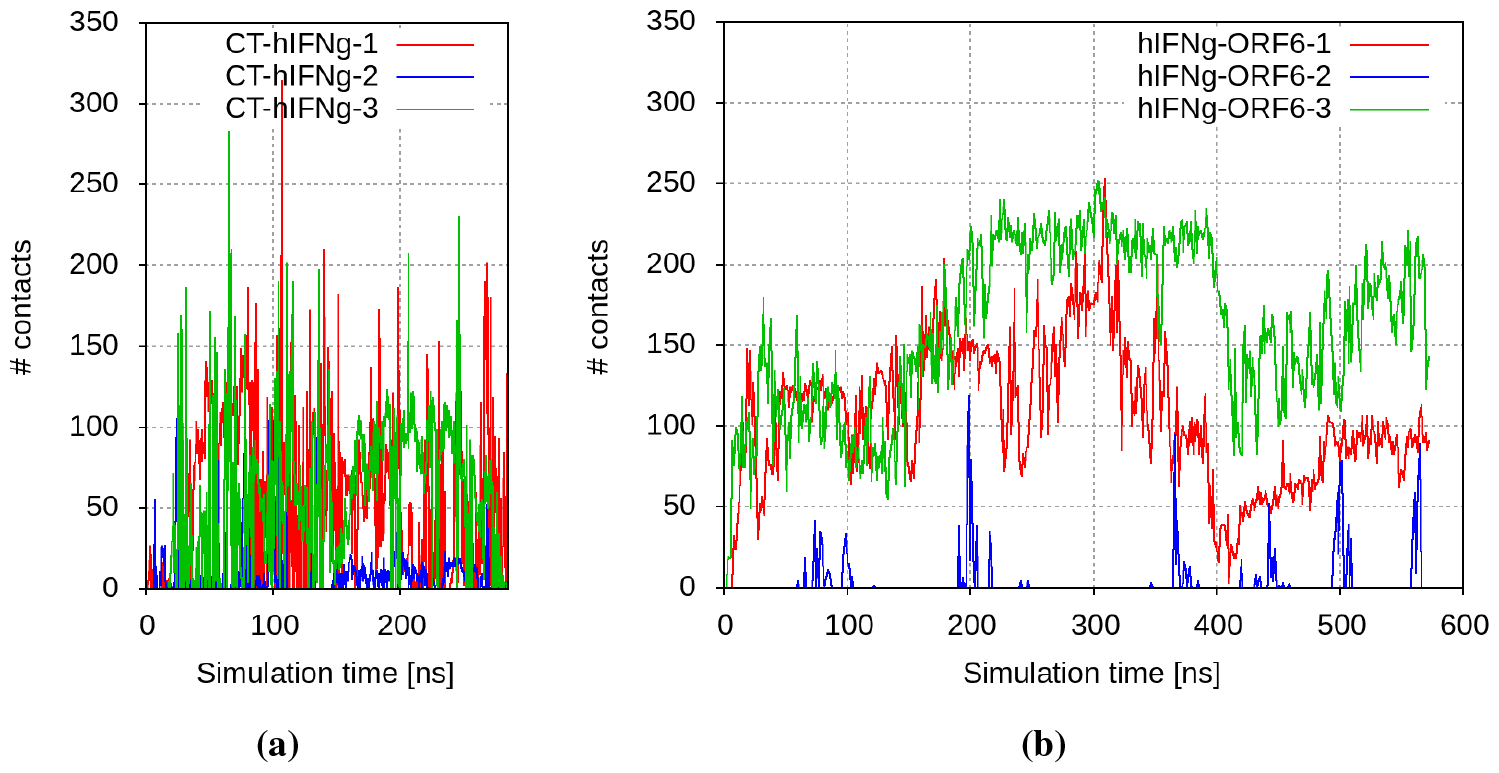
Number of contacts between (a) ORF6 and the CT-hIFN*γ* peptides; and (b) hIFN*γ* and the ORF6 proteins.

**Figure S3.**
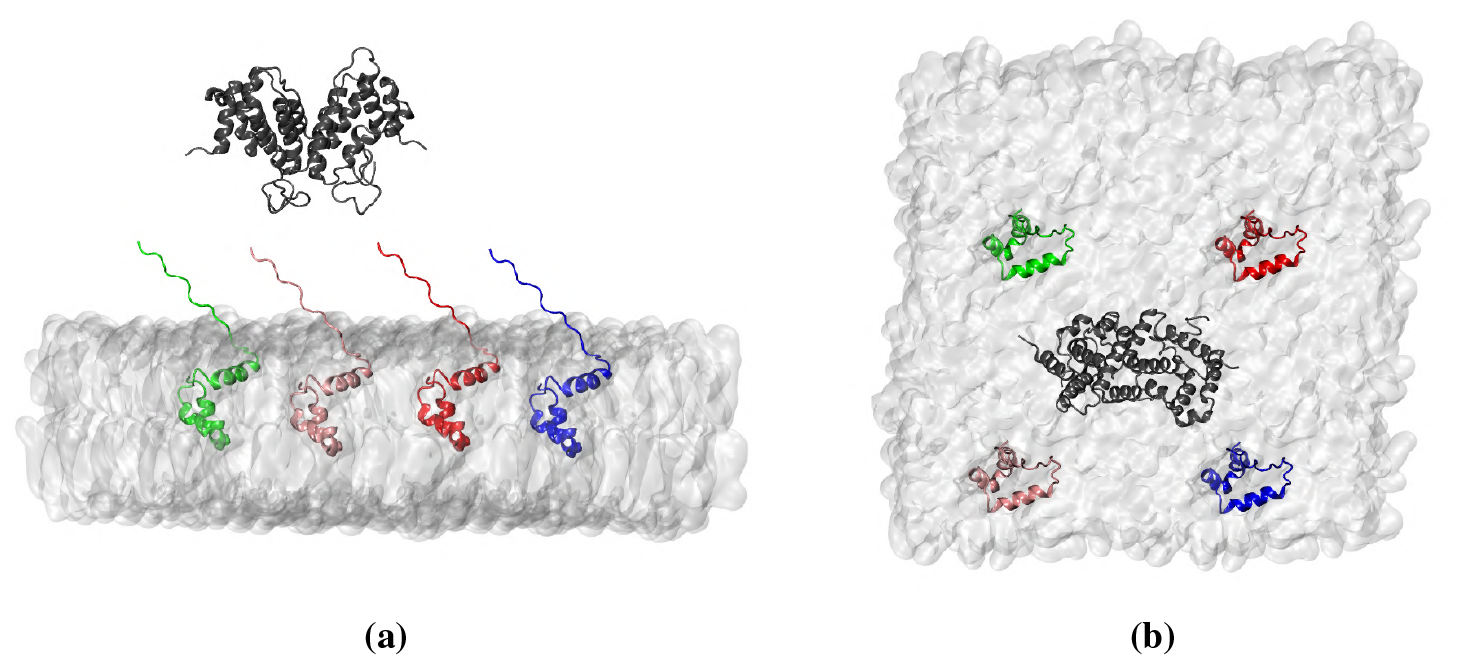
(a side view and (b) top vie of the input configuration of a full-length hIFN*γ* homdimer, placed above 4 ORF6 proteins, embedded in a model ER membrane. hIFN*γ* is depicted in dark gray, ORF6-1 to ORF6-4 are shown respectively in red, blue, green, and pink. The ER membrane is in light gray surface representation.

**Figure S4.**
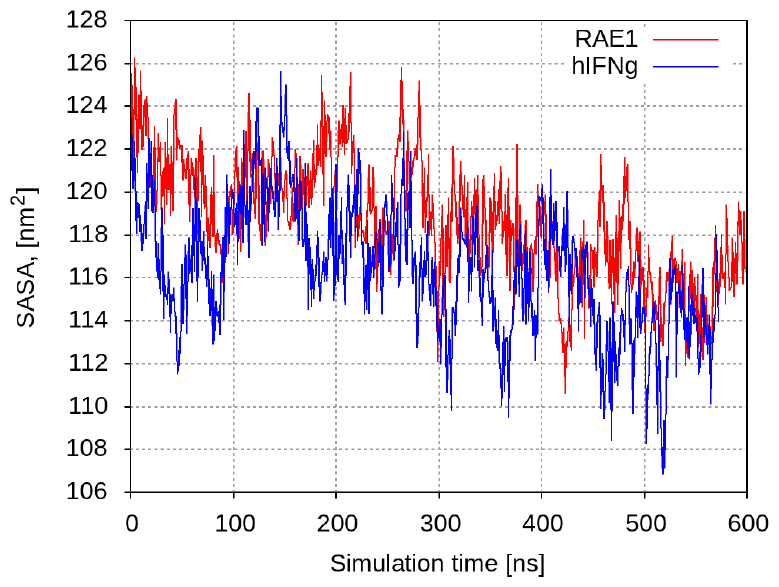
Evolution of the SASA of four ORF6 proteins when interction with RAE1 [9] or hIFN*γ*.

**Table S1.**
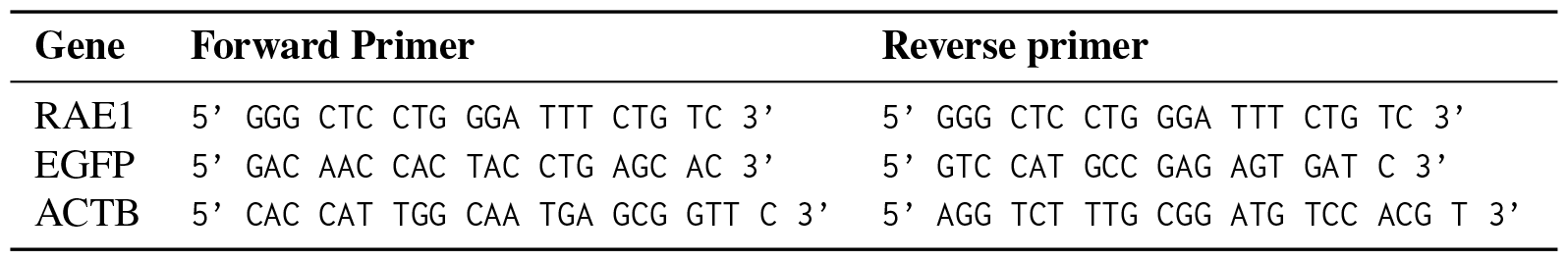
List of Primers

